# Binary search and set operations on compacted *k*-mer lists

**DOI:** 10.64898/2026.06.29.735436

**Authors:** Yoann Dufresne, Francesco Andreace

## Abstract

Sorted lists of elements are particularly good for computing set operations. A single scan of two lists is sufficient to materialize or count the results of the union, intersection, difference, and xor operators. In bioinformatics, only a few tools are designed to perform these operations on *k*-mers. A fast tool like KMC allows set operations at the cost of storing individual *k*-mers. In this paper, we introduce a novel way to represent sorted *k*-mers as a collection of recomposed super-*k*-mer sorted lists. We introduce the concept of virtual super-*k*-mer and show how to construct, query and perform set operations on sorted lists of virtual super-*k*-mers. In the implementation sklib, we demonstrate high throughput of the data structure for construction and set operations, while remaining competitive in query capabilities, within a controlled memory footprint (3.5–4.1*×* decrease in bits/element compared to KMC).

## 1 Introduction

The analysis of massive high-throughput sequencing collections is increasingly formulated as an indexing problem over sets of fixed-length substrings, termed *k*-mers. Representing datasets by their *k*-mer sets lets petabase-scale collections be stored in searchable structures, as demonstrated by MetaGraph [6] and Logan [4]. Analysis is no longer limited to asserting element presence: it increasingly requires operations over entire *k*-mer sets—intersections to identify shared content, differences to subtract contaminants, unions to incorporate new data, and symmetric differences to isolate dataset-specific variation— thus positioning *k*-mer indexing close to database and information-retrieval workloads where Boolean queries reduce to primitive set operations, as implemented in vizitig [5].

Sorted lists intrinsically support these operations: given two sorted *k*-mer sets, union, intersection, difference, and symmetric difference are computed by a single merge-like scan, and their cardinalities can be obtained without materializing the output. KMC [7] exploits this, but sorted lists store each element of a set *K* independently, so their space grows with *k*|*K*| despite adjacent genomic *k*-mers sharing *k* − 1 characters. Compact representations reduce this redundancy—filters and dictionaries [10,8,9] favor membership, BWT/FM-index structures [3,2,11] favor succinct queries—but they often require auxiliary indexes and hide the ordering needed for merge-based set algebra (Appendix A, Table 2).

We address this gap with virtual-super-*k*-mers: a representation of a *k*-mer set as a sorted list of recomposed super-*k*-mer-like objects that maintain an ordering over *k*-mers. Union, intersection, difference, and symmetric difference are thus still computed by sequential scans over compact virtual-super-*k*-mers rather than individual *k*-mers.

## 2 Super-*k*-mer representation

To exploit single-scan set operations and binary search, we must keep *k*-mers ordered in a list. We first outline the general idea of a compact sorted list of super-*k*-mers, then show how to construct them, how to query them and perform set operations on them, and finally we present optimizations that make the implementation efficient.

Three nested levels of partitioning appear throughout. A *bucket* is one of the 2^b^ independent lists the *k*-mer set is split into (Section 2). Inside a bucket, a *column* groups the *k*-mers whose minimizer occurs at one given position. A *record* is one stored entry of the final list, that is, one virtual-super-*k*-mer: it spans an interval of columns and carries one *k*-mer at each column of that interval. We reserve *list* for a sequence of records in sort order.

### 2D *k*-mer sorted list

A classical sorted list of *k*-mers cannot be directly compacted while preserving order; we therefore introduce a *2D sorting*. The real implementation never materializes it explicitly, but the whole method builds on this idea. A minimizer of size *m* is the smallest string of this size in a *k*-mer with respect to a given order (usually on hash values). We partition the *k*-mer set into columns according to minimizer position. There are exactly *k* − *m* + 1 possible positions for a minimizer inside a word of size *k*, so we construct *k* − *m* + 1 columns (Fig. 1). A *k*-mer whose minimizer starts at nucleotide *x* is sent to column *k* − *m* − *x* (the number of nucleotides following the minimizer). This minimizer position is the first dimension of sorting. The second dimension is the standard sort within each individual column, by a total order ≺ (alphabetical in our examples). We write *C*_i_ for the column of index *i*; the 2D list is thus the sequence of ≺-sorted columns *C*_0_, …, *C*_k*−*m_, and we write *u* ∈ *C*_i_ for a *k*-mer occurring at column *i*.

**Fig 1.**
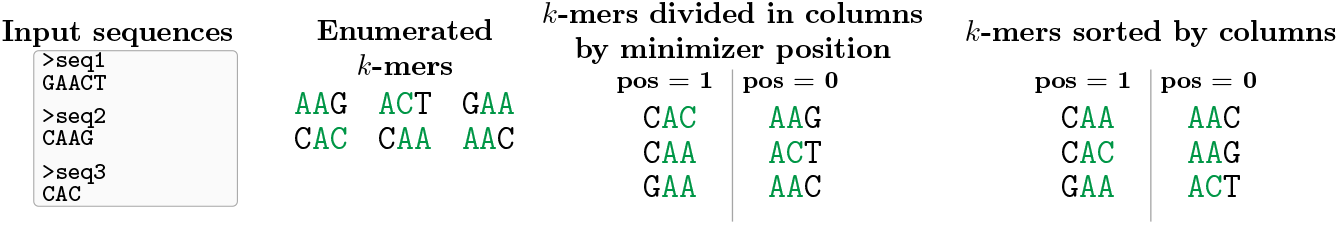
2D sorting for *k* = 3, *m* = 2. *k*-mers are enumerated from the sequences and their minimizers are highlighted. Then, *k*-mers are partitioned by *minimizer position* and column sorted. The labels pos = *x* give the start position of the minimizer inside the *k*-mer; the column index is *i* = *k* − *m* − *x*, so the left column (pos = 1) is *C*_0_ and the right one (pos = 0) is *C*_1_.

### Construction of super-*k*-mer lists

From the 2D list of the previous section, we want to produce a single 1D list of virtual-super-*k*-mers. We first distinguish classical super-*k*-mers from virtual-super-*k*-mers: the *k*-mers inside a classical super-*k*-mer are *collocated*, i.e. consecutive in the genome from which they are extracted, whereas virtual-super-*k*-mers drop this collocation requirement on their component *k*-mers. Virtual-super-*k*-mers are constructions needed for compaction, not objects carrying a biological meaning.

#### Algorithm 1

Building the sorted virtual-super-*k*-mer list.

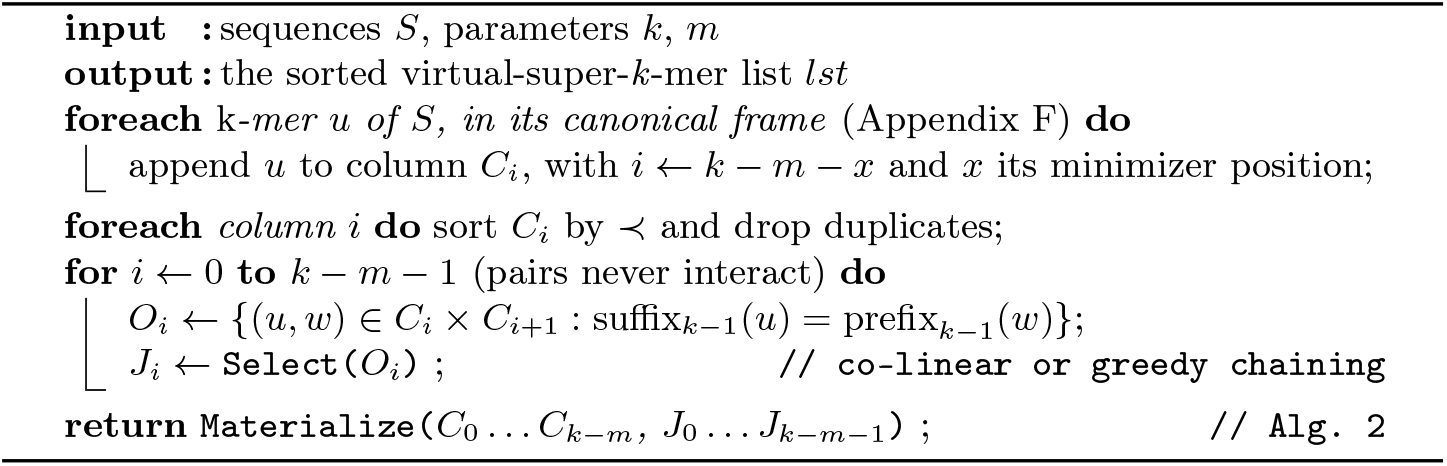

Formally, a virtual-super-*k*-mer *v* covers a non-empty interval of consecutive columns [*a*_v_, *b*_v_] ⊆ [0, *k* − *m*] and selects one *k*-mer per covered column, *v*_i_ ∈ *C*_i_ for each *i* ∈ [*a*_v_, *b*_v_], such that consecutive selections overlap on *k* − 1 nucleotides: suffix_k*−*1_(*v*_i_) = prefixk*−*1(*v*_i+1_) for all *i* ∈ [*a*_v_, *b*_v_ − 1]. Each index *i* ∈ [*a*_v_, *b*_v_] is a *position* of *v*, and *v*_i_ is the *k*-mer of *v* at that position; *v* spells a string of length *k* + (*b*_v_ − *a*_v_). A classical super-*k*-mer is the special case where the *k*-mers *v*_i_ are, in addition, collocated in a source sequence.

Our goal is to output a list *lst* of virtual-super-*k*-mers that respects the order ≺ of every column. Two virtual-super-*k*-mers can be compared only on the columns they share, which form an interval. Let [*a*_x_, *b*_x_] denote the span of *lst*[*x*], the list must satisfy:

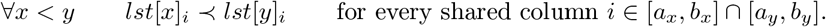

The quantifier thus ranges over the actual span shared by the two virtual-super-*k*-mers, not over the full range [0..*k* − *m*]. We build *lst* in two steps: we first *select*, between each pair of adjacent columns, a valid set of overlapping *k*-mers; we then *materialize* the resulting chains into the sorted list. Algorithm 1 states the construction as a whole.

### Selecting overlaps between adjacent columns

Two *k*-mers at adjacent columns *C*_j_ and *C*_j+1_ *overlap* if the last *k* − 1 nucleotides of a *k*-mer in *C*_j_ equal the first *k* − 1 of one in *C*_j+1_; such a pair of *k*-mers (*u, w*) ∈ *C*_j_ *× C*_j+1_ is a *candidate join* and could later be stored once, as a longer virtual-super-*k*-mer. A single *k*-mer may have several candidate joins (see Fig. 2). This step does not merge anything yet; it only *selects* which candidate joins will be used, subject to two constraints. First, the selection is a matching: each *k*-mer is kept in at most one join on each side, so selecting a join discards the remaining candidates of its two *k*-mers. Second, selected joins must not *cross*: two joins (*u, w*) and (*u*^*′*^, *w*^*′*^) cross when their *k*-mers appear in opposite orders in the two columns (*u* ≺ *u*^*′*^ but *w*^*′*^ ≺ *w*), and keeping both would invert the column order on one side. For instance, in Fig. 2, selecting the overlap between CAA and AAG (which would later form CAAG) forbids selecting the one between GAA and AAC (GAAC): column 0 orders CAAG before GAAC, whereas column 1 orders GAAC before CAAG, so no ordering of the two virtual-super-*k*-mers respects both columns.

**Fig 2.**
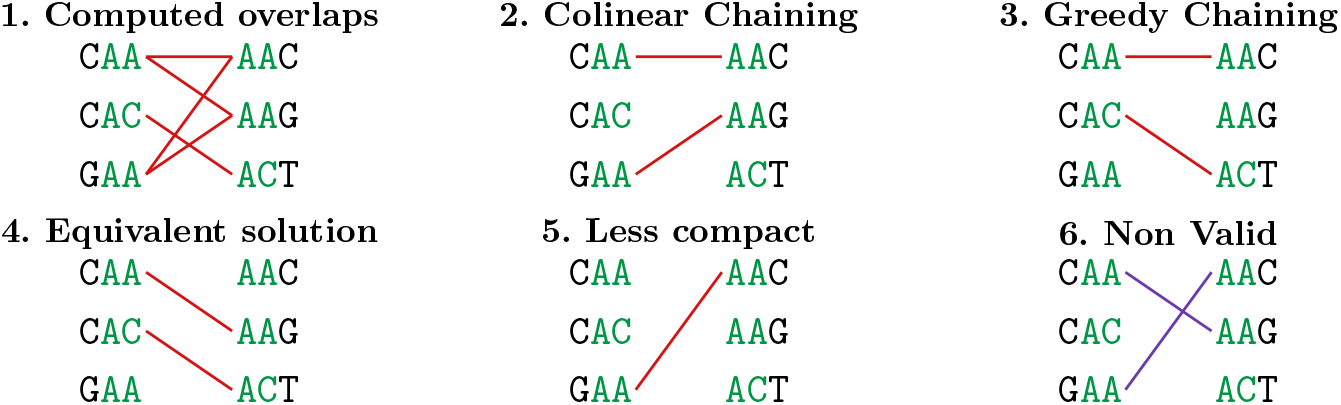
Co-linear chaining to select valid *k*-mer overlaps. Computed overlaps (1) are filtered to retain order-preserving chains between *k*-mer columns (2). Panels 3–5 show, for comparison, the greedy selection, an equivalent selection of the same size, and a valid but less compact one; panel 6 shows an invalid selection, whose crossing overlaps admit no order respecting both columns.

The sklib implementation provides two algorithms that select as many overlaps as possible between two columns. The first, *co-linear chaining* [1], is an *O*(*n* log *n*) dynamic program based on Fenwick trees that selects a maximum non-crossing set. The second is a greedy heuristic that always takes the first available overlap while sweeping the two lists from top to bottom; it is faster and, although it carries no guarantee, it selected exactly the same number of overlaps as the dynamic program on every genome we tried. It is therefore the default.

Selecting overlaps for one pair of columns has no effect on the selection for any other pair, since the *k*-mers keep their order in every column. Moreover, two equivalent (equally large) selections between the same pair yield the same *number* of virtual-super-*k*-mers; only their contents differ.

### Materializing the virtual-super-*k*-mer list

The selected joins chain *k*-mers across columns: each *k*-mer lies in one chain, and each chain spells one virtual-super-*k*-mer. These must be emitted as a sorted 1D list. We sweep all columns top-down in parallel, one pointer per column, and at each step emit the smallest virtual-super-*k*-mer whose *k*-mers are all pointed simultaneously; such virtual-super-*k*-mers occupy disjoint columns—so no column orders them—and ties break on the smallest interleaved encoding value (Section 2, Fig. 4). This yields a globally sorted, de-duplicated virtual-super-*k*-mer list (Algorithm 2). These records match enumerated super-*k*-mers in form (span ≤ 2*k* − *m*), but their boundaries come from the sort, not the reads. Still, the represented *k*-mer set is unchanged. sklib indexes canonical *k*-mers. Since a *k*-mer and its reverse complement carry their minimizer at mirrored positions, the canonical frame is chosen per *k*-mer rather than per record, so that both strands reach the same column; Appendix F states the rule.

#### Algorithm 2

Materializing the sorted virtual-super-*k*-mer list.

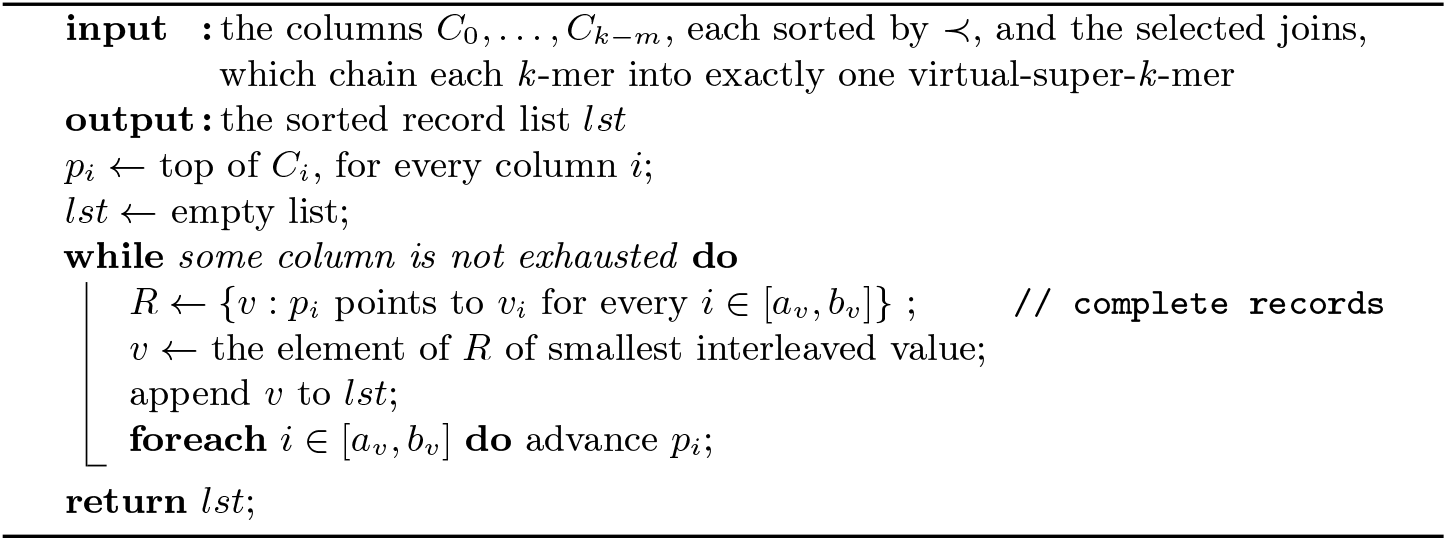

### Membership queries and set operations on sorted virtual-super-*k*-mer lists

Querying a *k*-mer is almost a plain binary search. At each step we mask the middle virtual-super-*k*-mer of the current range to expose the *k*-mer at the column given by the query’s minimizer position. When the masked virtual-super-*k*-mer carries no *k*-mer at that column, the algorithm scans linearly to the next virtual-super-*k*-mer that does, instead of the usual binary-search jump; the worst case scans the whole list but does not occur in practice. Fig. 3 runs it on an example.

**Fig 3.**
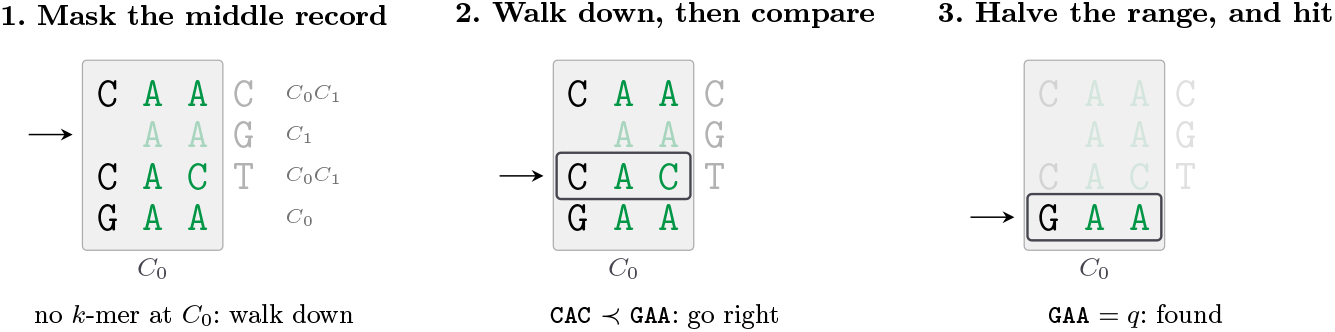
Masked binary search for the query GAA, on the list built from the *k*-mers of Fig. 1 (*k* = 3, *m* = 2). Each record is drawn at the columns it spans, which aligns every minimizer (green) on the same cells: one mask then exposes the *k*-mer of the queried column in every record at once. The middle record carries none at *C*_0_, so the search walks down to the next record that does (2), compares, and halves the range (3).

**Fig 4.**
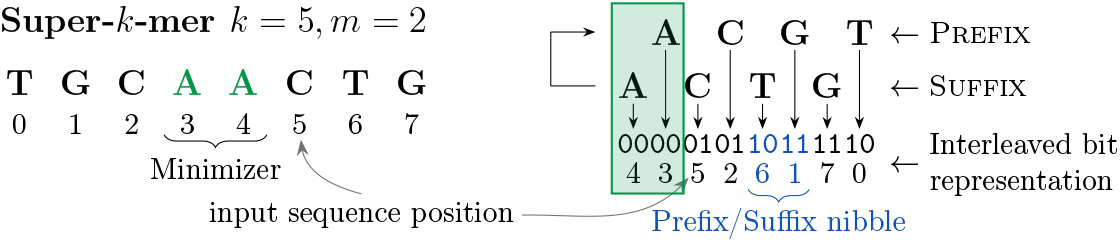
Interleaved, minimizer-centered encoding of a super-*k*-mer. The most significant bits contain the minimizer; prefix and suffix nucleotides are interleaved 4 bits at a time from the centre outward: a plain integer comparison realizes the centre-first sort key.

Since every *k*-mer has exactly one minimizer position, the columns partition the *k*-mer space: an operation between two lists decomposes into *k* − *m* + 1 independent two-way merges of sorted *k*-mer lists, one per column, and *k*-mers of different columns are never compared. The virtual-super-*k*-mers are only the storage layer of those lists—inside a column, a pointer moves from one record carrying a *k*-mer there to the next. Each step of a merge distinguishes three cases: the *k*-mer is present only in *A*, in both lists, or only in *B*; the pointer of the smaller *k*-mer advances, or both when they are equal. The four supported operations are four ways of selecting among those cases—keeping the second alone gives *A* ∩ *B*, keeping all three *A* ∪ *B*, the first alone *A \ B*, and the first and the third together the symmetric difference—so a single linear scan of the two lists produces any of them. An example is presented in Appendix C.

The result of an operation is itself a virtual-super-*k*-mer list: the kept *k*-mers are re-chained by the very procedure used for construction, so the representation is closed under its own operations and a result can feed the next one—re-chaining *A* ∪ *A* returns exactly the record count of *A*’s own index. That re-chaining is the largest phase of a materialized operation (half the wall time at one thread for a difference, three quarters for a union, against 22–42% for the merge itself), but this cost is intrinsic in materialization. Emitting one record per *k*-mer instead, which sklib also supports, still has to sort those *k*-mers, writes 8 to 15 times more, and came out slower in 395 of the 400 configurations we measured (Appendix D).

`sklib` implements two optimizations for this step, both of which fall out of the merge itself. The first lets several set operations be computed together: the three cases are selected independently of one another, so a single pass produces them all. The second computes cardinalities without materializing the result, making it several times faster (Appendix D), by counting the selected *k*-mers instead of collecting them, amounting to a single pass of memory reads.

### Optimization: minimizer-centered interleaving

Each nucleotide takes two bits (*A*=00, *C*=01, *T* =10, *G*=11, so the order ≺ realized by the implementation is *A* ≺ *C* ≺ *T* ≺ *G*; the examples of this paper use the alphabetical order instead, for readability, and any total order on *k*-mers is admissible), but instead of classical bit packing we use what we call an interleaved layout (see Fig. 4). Each super-*k*-mer is folded onto itself, with the folding point at the middle of the minimizer. This places the minimizer in the most significant bits, while the extreme nucleotides—present in only one *k*-mer of the super-*k*-mer—fall in the least significant bits. Shared nucleotides therefore have a strong influence on the integer value, whereas *k*-mer-specific nucleotides have little. This also ensures that two consecutive *k*-mers sharing a minimizer have similar values, and hence similar positions once their column is sorted, which allows more non-crossing overlaps between *k*-mers in adjacent columns. Two further optimizations make the implementation efficient: *bucketing* the records into 2^b^ independent lists (default *b*=12) for parallelism—legitimate because a join links only *k*-mers sharing the central (*k*−1)-mer and hence the same minimizer, so no chain ever crosses a bucket and each one is built, materialized and searched on its own—and *quotienting*, which drops the *b* bucket-implied bits per record since they are shared by all *k*-mers of a bucket. Both are detailed in Appendix E.

### 3 Experiments

We benchmark *E. coli*, yeast, *C. elegans*, and human chr1 (204.2 M) at *k* = 31 (*m* = 15) and *k* = 63 (*m* = 31), using *t* = 1 and *t* = 8 threads. The compared tools are listed in Appendix A (Table 2); all exact tools build canonical indexes, and KMC has no point-query mode. Each set-operation pair consists of a genome and a mutated copy of it, the substitution rate being derived from a target Jaccard index; the reported times are medians over those targets. The experimental setup, the mutation model, the tool versions and the build flags are given in Appendix I.

### Construction

On one thread sklib builds within 0.9–1.5*×* of KMC, the fastest constructor, and faster than every other tool: chr1 at *k*=31 takes 17.97 s, against 28.43 s for sshash and 73 to 236 s for the rest. It also scales better than KMC: with 8 threads the same index takes 3.52 s, a 5.1*×* speedup against KMC’s 4.6*×*, which puts sklib 1.8–2.3*×* ahead of KMC on *E. coli* and yeast, on par on *C. elegans*, and 1.1–1.4*×* behind on chr1. It uses far less memory: 177 MB peak RSS versus 735 MB for sshash and 1.8–9.0 GB for the others. Its space is intermediate—16–18 bits/*k*-mer, against 64–66 for KMC and 57–65 for cbl, yet above membership-oriented indexes (2–12 bits/*k*-mer)—thus much smaller than explicit sorted *k*-mer databases while retaining native set operations (full per-tool table in Appendix H). A record carries 7.9 to 9.0 *k*-mers at *k*=31 and 16.4 to 17.0 at *k*=63, about half of the *k* − *m* + 1 maximum in both cases; the space per *k*-mer follows directly from that ratio (Appendix G).

### Membership queries and set operations

Membership queries, though not sklib’s primary target, are competitive: on chr1 at *k*=31 sklib reaches 0.480 M*k*-mer/s against 0.347 for sbwtrs, with sshash—capped at *k* ≤ 31—still ahead at 0.633. From *k*=31 on sklib is 1.4 to 1.9*×* faster than sbwtrs, its throughput falling more slowly with *k* (0.323 against 0.181 M*k*-mer/s at *k*=63), and it answers absent *k*-mers as fast as present ones (full results in the repository). We compare exact set-operation tools to test whether virtual super-*k*-mers preserve merge-based performance under compaction. Table 1 reports results of the union in the case it materializes the whole result, which represents the operation on which sklib performs worst: there it stays within 1.5*×* of KMC at *k* = 31 and matches or beats it at *k* = 63, while cbl is 2.5–4.2*×* slower and sbwtrs/fmsi are 40–100*×* slower or limited to the smaller inputs. On the three other operations sklib is 1.2–4*×* faster than KMC, because its cost follows the size of the result where KMC’s does not: on chr1 it ranges from 18.3 s for the union of two disjoint sets down to 0.97 s for a difference between near-identical ones, a 19*×* range against 1.4*×* for KMC (Appendix B). sklib also becomes faster than KMC with threads (2.3*×* at *k* = 31, 3.3*×* at *k* = 63, *t* = 8; Table 3) while using far less memory: 24 MB versus 2.5 GB for KMC and 22.4 GB for cbl at *k* = 31, *t* = 1.

**Table 1.** Set operations: time (s) for the union *A* ∪ *B*, single-threaded (*t* = 1), median over seven Jaccard targets. The union is reported because it materializes the whole result and is the operation on which sklib does worst; sklib also supports ∩, *\* and the symmetric difference. Every operation at every target is given in Appendix B (Table 4). “–” marks unsupported or capped configurations. Thread scaling and memory on chr1 are reported in Appendix B (Table 3).

| Tool | $k = 31$ | | | | $k = 63$ | | | |
| --- | --- | --- | --- | --- | --- | --- | --- | --- |
|  | *E. coli* | Yeast | *C. elegans* | Chr1 | *E. coli* | Yeast | *C. elegans* | Chr1 |
| sklib | 0.257 | 0.681 | 5.667 | 12.94 | 0.282 | 0.700 | 5.841 | 13.09 |
| KMC | 0.229 | 0.991 | 5.016 | 8.629 | 0.448 | 1.008 | 7.370 | 13.04 |
| cbl | 0.711 | 1.697 | 24.01 | 50.19 | — | — | — | — |
| sbwtrs | 10.64 | 26.64 | — | — | — | 58.90 | — | — |
| fmsi | 20.56 | 55.93 | — | — | 23.67 | 70.85 | — | — |

## 4 Conclusion and future work

We presented a compact representation of sorted *k*-mer sets as an ordered list of virtual super-*k*-mers. The key idea is to compact overlapping *k*-mers into minimizer-based objects while preserving the sorted-list structure that enables efficient merge-based set operations. Our experiments show that our implementation, sklib, occupies an intermediate point between highly compact membership indexes and explicit sorted *k*-mer databases: it uses substantially fewer bits per element than KMC, remains competitive for membership queries, and is faster than KMC on set operations—on three of the four at one thread, on all four with eight threads—with much lower memory usage. These results suggest that virtual super-*k*-mers provide a practical way to compact *k*-mer sets without losing the advantages of sorted representations. In the near future we will apply these set operations to the de novo construction of bacterial pangenomes, and to improving database operations on *k*-mers together with the authors of Vizitig [5].

## Supporting information

All supplementary

## Acknowledgments

F.A. was supported by ANR Full-RNA and Inception grants (ANR-22-CE45-0007, PIA/ANR16-CONV-0005). This project has received funding from the European Union’s Horizon 2020 research and innovation programme under the Marie Skłodowska-Curie grants agreements No. 872539 and 956229.

