## Supplementary material for "Binary search and set operations on compacted *k*-mer lists": All supplementary

### A Compared tools

Table 2 details the tools benchmarked in Section 3.

**Table 2.** Tools compared in Section 3. "Set-ops" marks native set operations between two $k$-mer sets; "Threaded" lists the workloads each tool parallelizes (C: construction, Q: queries, S: set operations). $^{\star}$bqf is an approximate filter (false positives), shown for reference only.

| Tool | Representation | $k$ range | Query | Set-ops | Threaded |
|---|---|---|---|---|---|
| sklib | sorted virtual super-$k$-mer list | $\leq 127$ | ✓ | ✓ | C, Q, S |
| KMC [7] | $k$-mer counter / on-disk DB | all | – | ✓ | C, S |
| sshash [10] | minimal perfect hash | $\leq 31$ | ✓ | – | – |
| sbwt [3] | spectral BWT (C++) | $\leq 32$ | ✓ | – | – |
| sbwtrs [2] | spectral BWT (Rust) | $\leq 255$ | ✓ | ✓ | C, S |
| cbl [9] | Conway–Bromage–Lyndon | odd $\leq 59$ | ✓ | ✓ | – |
| fmsi [11] | masked superstring (FM-index) | $\leq 127$ | ✓ | ✓ | – |
| bqf$^{\star}$ [8] | quotient filter (approximate) | any | ✓ | – | – |

### B Set operations: thread scaling on chr1

Table 3 reports thread scaling and peak memory on chr1, complementing the single-threaded results of Section 3.

**Table 3.** Set operations on chr1: thread scaling and memory for sklib and KMC, on the union, as in Table 1.

| | time $k = 31$ | | time $k = 63$ | | RAM $k = 31$ (MB) | |
|---|---|---|---|---|---|---|
| Tool | $t$=1 | $t$=8 | $t$=1 | $t$=8 | $t$=1 | $t$=8 |
| sklib | 12.94 | 3.21 | 13.09 | 3.60 | 24 | 107 |
| KMC | 8.63 | 7.41 | 13.04 | 11.76 | 2525 | 2527 |

*Cost across operations and overlap.* Table 4 shows the values underlying the median in Table 1 on chr1. sklib's time follows the number of $k$-mers the result holds: across these targets the intersection grows from 0 to 204 M $k$-mers, the union shrinks from 435 to 204 M, and a difference empties from 204 M to none. KMC takes 7.6 to 10.8 s because it re-reads and re-dumps its database either way; cbl is flat (1.7× across the grid). The last line of the table is the ratio between the most and the least expensive target of each column, which reaches 47× for sklib once the two empty differences at $J = 1$ are counted. Results are consistent with the other three smaller datasets.

**Table 4.** Set operations on chr1 ($k = 31$, $t = 1$): time (s) for each operation at each Jaccard target, sklib against KMC. The last line is the ratio between the most and the least expensive target of that operation—what a median over the targets summarizes.

| | $A \cap B$ | | $A \cup B$ | | $A \setminus B$ | | $B \setminus A$ | |
|---|---|---|---|---|---|---|---|---|
| $J$ | sklib | KMC | sklib | KMC | sklib | KMC | sklib | KMC |
| 0 | 3.32 | 10.50 | 18.34 | 10.74 | 10.59 | 10.77 | 10.85 | 10.39 |
| 0.1 | 5.09 | 9.54 | 17.75 | 9.28 | 9.04 | 8.95 | 9.88 | 9.10 |
| 0.3 | 6.99 | 8.83 | 15.35 | 8.74 | 6.88 | 8.43 | 7.47 | 8.64 |
| 0.5 | 7.72 | 8.87 | 12.94 | 8.49 | 4.57 | 8.31 | 4.81 | 8.21 |
| 0.7 | 7.84 | 9.10 | 11.08 | 8.51 | 2.69 | 7.96 | 2.73 | 7.90 |
| 0.9 | 8.36 | 8.80 | 9.05 | 8.44 | 0.97 | 7.78 | 1.03 | 7.81 |
| 1 | 8.29 | 8.67 | 8.16 | 8.63 | 0.22 | 7.65 | 0.23 | 7.61 |
| spread | 2.5× | 1.2× | 2.2× | 1.3× | 47× | 1.4× | 47× | 1.4× |

### C Set operation merging example

Fig. 5 shows an example of the merge-like scan of two lists computing $A \cap B$. Consider two lists $A$ and $B$, each made up of two columns of $k$-mers. The set operation scan starts by placing pointers on top of each column to keep track of $k$-mers to be compared. To make comparison clear, the left column of each list is pointed, in the figure, by an arrow, while the right is pointed by a small circle. At the beginning, list $A$ exposes `CAA` and `AAC`, while list $B$ exposes `AAA` then `AAT`: `CAA` of $A$ is compared to `AAA` of $B$ and `AAC` of $A$ is compared to `AAT` of $B$. Therefore as `AAA`≺`CAA`, `AAA` is discarded and the little arrow advances in list $B$ to the next virtual-super-$k$-mer in the column. In the same way, `AAC`≺`AAT`, so `AAC` is discarded and the little circle advances in list $A$ to the next virtual-super-$k$-mer with a $k$-mer in the right column. Comparison by column continues, and in the right column both list $A$ and $B$ expose `AAG`: as the two $k$-mers are equal, `AAG` is emitted and the little circle pointer advances in both lists. Comparison is continued until both lists maintain elements in the same column, else the remaining elements in the list are discarded (or emitted for other set operations). Replacing the emit rule turns the same single scan into $\cup$, $A \setminus B$, or the symmetric difference.

### D Cost of materializing a set operation

Per bucket, a materialized set operation reads the two operand buckets, merges them column by column to decide which $k$-mers the result holds, orders the kept $k$-mers into the output structure, and writes it. Only the third phase distinguishes the two ways of emitting a result: by default sklib hands the kept $k$-mers to the construction routine, which chains them into virtual-super-$k$-mers, whereas `--no-compact` sorts them and keeps one record per $k$-mer. The `*_size` variants stop after the merge and write nothing, so they time the merge alone. The three modes were run on the grid of Section 3 (four operations, seven Jaccard targets,

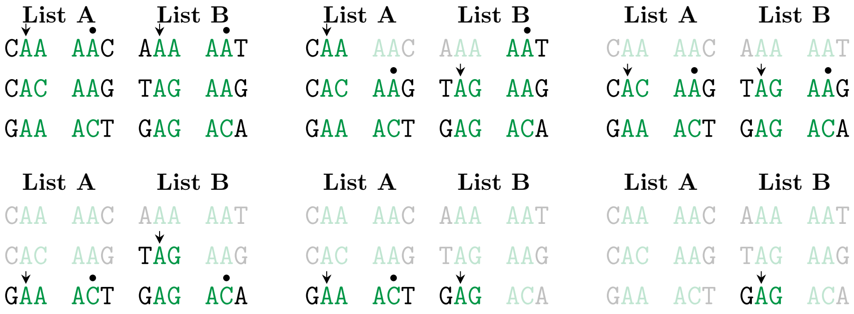


**Fig. 5.** Set operation scan of the virtual super-$k$-mer sorted list visualized as 2D sorting of $k$-mers.

medians of three runs), with the two differences at $J = 1$ being empty and therefore excluded.

Table 5 shows the single-thread wall time. The chaining is the largest phase of every operation, and its share follows how much of the operands the result keeps: 49–51% for a difference, 63–64% for an intersection, 73–74% for a union, where the merge step takes only 22% of the total time. In the uncompacted mode, at $k = 31$, sorting the kept $k$-mers costs exactly the same time as chaining (0.98–1.01×), and at $k = 63$, where $k - m + 1 = 33$ columns leave a global sort much more to order, sorting is the more expensive of the two (1.43–1.54×). Chaining also divides the bytes written by 4.7 to 11.9.

**Table 5.** Materialized operation elapsed time breakdown, at $t = 1$, one line per operation (medians over the four datasets and the seven Jaccard targets; dataset difference is within 2 points). The four shares are of the per-bucket wall time reported by the library's own phase timers. The last two columns give `--no-compact` costs relative to default: sorting the kept $k$-mers instead of chaining them, and writing the result.

| | | share of the per-bucket time | | | | `--no-compact` / default | |
|---|---|---|---|---|---|---|---|
| Operation | $k$ | read | merge | chain | write | order | write |
| $A \cap B$ | 31 | 2.7% | 32.2% | 62.7% | 2.1% | 1.01× | 5.4× |
| | 63 | 2.6% | 29.8% | 64.2% | 2.2% | 1.51× | 10.1× |
| $A \cup B$ | 31 | 1.6% | 22.2% | 74.0% | 2.4% | 1.01× | 6.0× |
| | 63 | 1.6% | 22.3% | 73.3% | 2.3% | 1.54× | 11.9× |
| $A \setminus B$ | 31 | 3.8% | 41.7% | 49.3% | 2.0% | 0.98× | 4.7× |
| | 63 | 4.0% | 36.5% | 56.0% | 2.1% | 1.44× | 8.5× |
| $B \setminus A$ | 31 | 3.7% | 40.9% | 51.0% | 2.0% | 0.98× | 4.8× |
| | 63 | 3.9% | 35.9% | 56.8% | 2.1% | 1.43× | 9.1× |

Table 6 gives the end-to-end times on chr1, the largest dataset. Materializing a result costs 2.4 to 7.2× a cardinality-only pass, ordered by how much the result keeps (a difference is at the low end, a union at the high one) which is the cost of turning a count into a list. Skipping the chaining into virtual-super-$k$-mers makes the uncompacted mode slower for every operation, by 1.1–2.0× at one thread and 1.2–2.2× at eight, and over the whole test grid it performed worse in 395 of the 400 configurations measured; the five exceptions are runs below 80 ms with a small result, the largest gain among them being 11%. Its output also holds one record per $k$-mer, that is exactly 144 bits/$k$-mer at $k = 31$ and 272 at $k = 63$, against the 17.1–20.6 of the compacted result. The three smaller datasets behave the same up to scale: per operation, the uncompacted mode costs 1.09–2.51× the compacted one and materializing costs 2.1–8.6× a count.

**Table 6.** Set operations on chr1 in the three modes: time (s), one line per operation (medians over the seven Jaccard targets). "count" materializes nothing and returns the cardinality; "uncomp." writes one record per result $k$-mer; "comp." re-chains the result into virtual-super-$k$-mers. The last column is the space of the compacted result, whereas the uncompacted one uses 144 bits/$k$-mer ($k = 31$) or 272 ($k = 63$).

| | | $t = 1$ | | | $t = 8$ | | | |
|---|---|---|---|---|---|---|---|---|
| Operation | $k$ | count | uncomp. | comp. | count | uncomp. | comp. | bits/$k$-mer |
| $A \cap B$ | 31 | 1.890 | 9.244 | 7.781 | 0.292 | 2.271 | 1.744 | 19.2 |
| | 63 | 1.614 | 14.587 | 8.260 | 0.284 | 4.363 | 1.970 | 17.7 |
| $A \cup B$ | 31 | 2.109 | 14.546 | 12.943 | 0.385 | 4.075 | 3.212 | 18.7 |
| | 63 | 1.823 | 26.059 | 13.086 | 0.408 | 7.573 | 3.595 | 17.1 |
| $A \setminus B$ | 31 | 2.429 | 6.439 | 5.726 | 0.482 | 1.619 | 1.310 | 20.6 |
| | 63 | 2.143 | 8.803 | 5.611 | 0.541 | 3.089 | 1.407 | 19.5 |
| $B \setminus A$ | 31 | 2.427 | 6.882 | 6.136 | 0.489 | 1.749 | 1.383 | 20.3 |
| | 63 | 2.173 | 8.778 | 5.924 | 0.539 | 2.859 | 1.469 | 19.2 |

Re-chaining $A \cup A$ returns exactly the record count of $A$'s own index, on all eight dataset/$k$ configurations, proving a chain of operations accumulates no loss. The uncompacted and compacted results were also checked to represent the same set, to answer the same queries identically, and to both feed a further set operation. The scripts, the raw table and the verification are in the repository.

### E Implementation optimizations: bucketing and quotienting

These two optimizations complement the minimizer-centered interleaving of Section 2; they leave every operation described in the main text unchanged.

*Bucketing.* sklib holds not one $k$-mer list but $2^b$ lists, the *buckets* ($b$ is a user parameter, default 12). The bucket of a $k$-mer is chosen by a hash function with good distribution properties: we hash every $m$-mer of the $k$-mer and keep the smallest hash (the minimizer). The record stores that hash bit-reversed, and the bucket is read off the $b$ highest bits of the reversed value, that is, the $b$ lowest bits of the hash itself. The reversal serves two purposes. First, a minimum search biases the high bits of the hash towards zero, so bucketing on them directly would leave most buckets nearly empty, whereas the low bits stay uniform. Second, it places the bucket identifier at the top of the record, hence as a prefix of the sort key: concatenating the buckets then yields a globally sorted list, and those $b$ bits become redundant in every record, which is what quotienting exploits. So that the $k$-mers can still be enumerated, the hash function is reversible; we use an xor-shift from the hash-prospector project [12]. Every operation described in the main text is unchanged, except that it first selects the relevant bucket before performing the list operations.

*Quotienting.* Because each $k$-mer is sent to a deterministic bucket based on $b$ of its bits, those bits need not be stored: the bucket address holds them implicitly, since they are shared by all the $k$-mers it contains. This saves $b$ bits per record, i.e. 12 bits with the default $b$=12.

### F Canonical $k$-mers and minimizer position

sklib indexes canonical $k$-mers: a $k$-mer and its reverse complement are treated as a single element, stored in one canonical orientation. This interacts with the 2D sorting because the minimizer of a $k$-mer and that of its reverse complement occur at mirrored positions within the word. Choosing the orientation only at the level of the full record is therefore insufficient. When the minimal $m$-mer is self-reverse-complementary or occurs multiple times within the window, the leftmost-occurrence rule used while sliding along a sequence is not mirror-symmetric: forward and reverse-complement scans select mirrored occurrences. As a result, the same $k$-mer can be assigned to different columns depending on the strand from which it is read, causing queries to miss one of the representations.

Fig. 6 shows such a $k$-mer. sklib therefore fixes the frame per $k$-mer rather than per record, with a rule that is symmetric by construction: among the occurrences of the minimal $m$-mer, keep those of minimal rank, break ties by taking the most central occurrence, and break remaining ties by the smaller canonical interleaved value. Both strands then derive the same frame from their own nucleotides, so a $k$-mer built in the context of a sequence and the same $k$-mer queried in isolation reach the same column. Records whose framing is ambiguous are split into per-$k$-mer pieces at construction time. As a consequence the index is exact for every value of $m$.

### G Record layout and space accounting

A record is stored in a fixed-width slot: the two halves of the interleaved pair, at an integer width selected at run time among 4, 8, 16 and 32 bytes, plus one byte for the prefix length and one for the suffix length. A record spans at most $2k - m$ nucleotides, so its prefix and its suffix are each at most $k - m \leq 127$ and one byte apiece suffices. Inside a bucket, the pairs and the length bytes are held in two separate arrays, because the binary search, the scan to the closest valid record and the set-operation merge each touch only one of the two. Quotienting

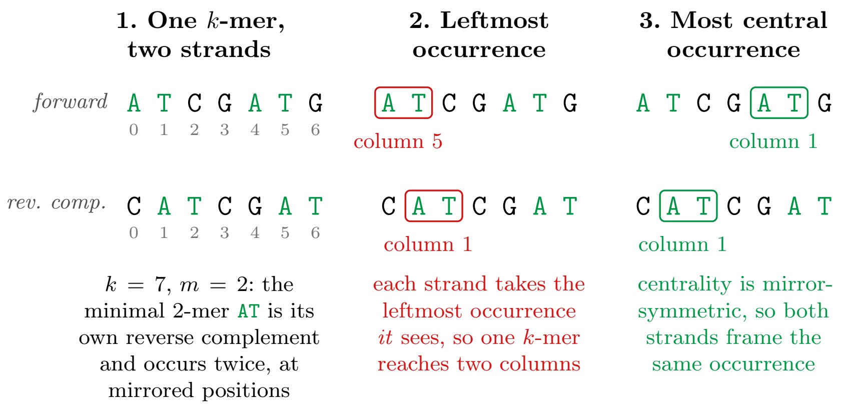


**Fig. 6.** Why the frame is fixed per $k$-mer. The $k$-mer `ATCGATG` and its reverse complement `CATCGAT` are one canonical element; its minimal 2-mer `AT` is its own reverse complement and occurs twice, at mirrored positions. Framing on the leftmost occurrence seen by each scan sends that single $k$-mer to two different columns; framing on the most central occurrence, a criterion unchanged by mirroring, sends both strands to the same one.

removes the $b$ leading bits of every record (Appendix E), which is what allows a narrower integer width to be selected.

The index size is therefore the number of records times the slot size, plus a directory of two 64-bit words per bucket (its lowest minimizer value and its record count), and the space per $k$-mer is entirely determined by the number of records per $k$-mer. At $k = 31$, $m = 15$ the selected width is 8 bytes and a record occupies 18; at $k = 63$, $m = 31$ it is 16 bytes and 34. Table 7 reports the compaction measured on the four datasets: a record carries close to half of the $k - m + 1$ $k$-mers it could hold, at both values of $k$, and these counts reproduce the space of Table 8 exactly. Since the greedy heuristic selects as many overlaps as the co-linear program, they are also the counts of a maximum non-crossing selection.

**Table 7.** Compaction achieved by virtual-super-$k$-mers, at one thread. “max” is the largest number of $k$-mers a record can hold, $k - m + 1$.

| | $k = 31$, $m = 15$ (max 17) | | $k = 63$, $m = 31$ (max 33) | |
|---|---|---|---|---|
| Dataset | records/$k$-mer | $k$-mers/record | records/$k$-mer | $k$-mers/record |
| *E. coli* | 0.112 | 8.96 | 0.059 | 17.00 |
| Yeast | 0.112 | 8.91 | 0.059 | 16.95 |
| *C. elegans* | 0.123 | 8.16 | 0.059 | 16.83 |
| Chr1 | 0.126 | 7.91 | 0.061 | 16.40 |

### H Construction space: bits per $k$-mer

Table 8 reports the space usage of all benchmarked tools, complementing the summary given in Section 3.

**Table 8.** Space usage in bits per $k$-mer for all tools at $k = 31$, sorted by usage on *E. coli*. $^{\star}$bqf is an approximate filter, shown for reference only.

| Tool | *E. coli* | Yeast | *C. elegans* | Chr1 |
|---|---|---|---|---|
| fmsi | 2.13 | 2.15 | 2.20 | 2.25 |
| sshash | 7.12 | 7.34 | 8.33 | 9.67 |
| sbwtrs | 11.84 | 10.73 | 10.09 | 10.04 |
| sbwt | 12.34 | 11.23 | 10.59 | 10.54 |
| **sklib** | **16.19** | **16.21** | **17.66** | **18.20** |
| bqf$^{\star}$ | 38.68 | 29.02 | 24.27 | 21.04 |
| cbl | 64.90 | 60.20 | 57.34 | 56.76 |
| KMC | 66.30 | 64.91 | 64.11 | 64.05 |

### I Experimental Setup

Experiments were run on an Intel Core Ultra 7 165H machine with 62 GiB RAM and Linux 6.17. Wall time and peak RSS were measured with `/usr/bin/time -v`; we report median time over three runs and maximum RSS. For the threaded tools the first rep of each configuration is a cold-page-cache run: the median discards it, so the tables show warm steady-state. We benchmark *E. coli* (4.6 MB, 4.5 M distinct $k$-mers), yeast (12 MB, 11.6 M), *C. elegans* (98 MB, 94.0 M), and human chr1 from CHM13 (224 MB, 204.2 M) at $k = 31$ ($m = 15$) and $k = 63$ ($m = 31$), using $t = 1$ and $t = 8$ threads. Full tables and scripts are in the repository; Appendix J maps each result to its file.

*Set-operation operands.* Each pair $(A, B)$ is a genome and a mutated copy of it: substitutions are applied at a rate $r$ derived from a target Jaccard index $J$ by $r = 1 - (2J/(1 + J))^{1/k}$, so that the two sets reach the intended similarity. The targets are $J \in \{0, 0.1, 0.3, 0.5, 0.7, 0.9, 1\}$, and the Jaccard index actually obtained is recomputed from the built lists and cross-checked against KMC. The times of Table 1 are medians over the four operations and over these targets, which is why they cover the whole similarity range rather than one arbitrary pair. Pairs of real genomes (*E. coli* K-12 against Sakai, UTI89 and IAI39; chr20 against chr21) are also provided in the repository.

*Tool versions and build flags.* Table 9 records what was built and how. Every tool was compiled through its own project's release configuration, so the flags differ between them; the two asymmetries worth stating are that sklib and bqf are the only ones compiled for the host CPU—the Rust tools in particular receive no `target-cpu` override—and that fmsi defaults to `-O2`. A portable sklib target (`-march=x86-64`) is available for reproduction on other machines.

**Table 9.** Version and build of every benchmarked tool. Flags are those the project's own release configuration enables, not flags we chose. cbl is compiled per $k$ (one binary for each value).

| Tool | Version / commit | Lang. | Optimization flags |
|---|---|---|---|
| sklib | v0.15.0 | C++ | `-O3 -march=native`, LTO |
| KMC | 3.2.4 | C++ | official release binary |
| sshash | `3e4f90b` | C++ | `-O3 -mbmi2 -mavx2` |
| sbwt | `b6e6830` | C++ | `-O3`, `MAX_KMER_LENGTH=32` |
| sbwtrs | `7c5fcc0` | Rust | `cargo build --release` |
| cbl | `986d57e` | Rust | `cargo +nightly build --release` |
| fmsi | `39c71a1` | C++ | `-O2` |
| bqf | `c9ff280` | C++ | `-Ofast -march=native -mtune=native` |

### J Raw data and reproduction

Every metric shown in this manuscript has been computed and stored in a table in the sklib repository, `https://github.com/yoann-dufresne/sklib` (tool version v0.15.0; data as of commit `f10164f`). Each CSV holds one row per measurement (tool version, host, dataset, $k$, $m$, threads, and the raw timing). Table 10 maps each result to its file and to the script that produced it.

**Table 10.** Source of each result reported in this manuscript. Data files are relative to `benchmark/results/` in the repository, scripts to `benchmark/scripts/`; `runs/.../` abbreviates `runs/full_run_2026-06/data/`, the campaign that produced every competitor figure.

| Reported in | Data file | Script |
|---|---|---|
| Tables 1, 3, 4; Appendix D (sklib) | `reference/setops_rechain.csv` | `setop_rechain.sh` |
| Tables 1, 3, 4 (other tools) | `runs/.../setop.csv` | `setop.sh` |
| Section 3, construction and space; Tables 7, 8 (sklib) | `reference/construct_v0.15.0.csv` | `construct.sh` |
| Table 8 (other tools) | `runs/.../construct.csv` | `construct.sh` |
| Section 3, membership queries (sklib) | `reference/query_single_v0.15.0.csv` | `query_single.sh` |
| Section 3, membership queries (other tools) | `runs/.../query_single.csv` | `query_single.sh` |
| Appendix D, correctness of the two emission modes | — (checks only, no table) | `verify/rechain_verif.sh` |
| Section 2, greedy versus co-linear chaining | `journals/GREEDY_DEFAULT.md` | `verify/greedy_chaining_verif.sh` |

The operands of the set-operation grid are generated by `scripts/mutate.py` at the substitution rate that `scripts/lib.sh` derives from each Jaccard target, and the raw rows are aggregated into the medians of Tables 4–6 by `scripts/rechain_report.py`. Re-running a campaign is one command, for instance

```
DATASETS="ecoli yeast celegans chr1" KM="31,15 63,31"
    bash benchmark/scripts/setop_rechain.sh
```

which rebuilds the indexes it needs, skips the rows already present in the CSV, and appends the new ones. Two write-ups in `results/journals/` record how the campaigns behind Appendix D and behind the query figures were run and what they found: `SETOP_RECHAIN.md` and `QUERY_V0150.md`.
